# Refining filtering criteria for accurate taxonomic classification of ancient metagenomic data

**DOI:** 10.1101/2025.03.31.646431

**Authors:** Nikolay Oskolkov

## Abstract

Taxonomic profiling is a key component of ancient metagenomic analysis, however it is also susceptible to false-positive identifications. In particular, classification tools from the Kraken family, such as Kraken2 and KrakenUniq, are highly sensitive to the choice of filtering options. To address this issue, various filtering approaches have been proposed. In this study, I conduct a comprehensive benchmarking of different filtering strategies using simulated microbial and environmental ancient metagenomic data. I evaluate these approaches based on the balance between sensitivity and specificity of ground truth reconstruction (F1-score), and propose an optimal thresholding strategy tailored to specific sequencing depths in ancient metagenomic datasets.

## Introduction

Metagenomic taxonomic profiling is a fundamental tool for studying both ancient and modern microbial communities, providing insights into past and present ecosystems. A variety of taxonomic classification tools, such as KrakenUniq [1], Kraken2 [2], MetaPhlan [3], and Centrifuge [4], have been developed to analyze metagenomic datasets. However, selecting the most appropriate classifier remains a subject of debate, particularly in the field of ancient metagenomics, where challenges such as low coverage (limited amount of DNA), short read length, DNA degradation and modern contamination complicate taxonomic assignments.

Several benchmarking studies have systematically compared different metagenomic taxonomic classification approaches, assessing their accuracy, sensitivity, and specificity across diverse datasets [5–7]. These studies highlight the strengths and limitations of various tools, emphasizing factors such as database completeness, algorithmic differences, and computational efficiency. However, a critical question remains: whether the true performance of classification algorithms themselves, or simply the impact of post-classification filtering strategies has been compared? Since different tools employ distinct filtering thresholds and confidence scoring mechanisms, variations in performance may often be attributed to differences in downstream processing rather than fundamental differences in classification algorithms.

Thus, a more nuanced approach is required when interpreting benchmarking results, one that considers not only the taxonomic classifiers but also the filtering strategies applied to their outputs. Understanding these distinctions is essential for optimizing metagenomic taxonomic profiling, particularly in the context of ancient DNA studies, where accurate classification is crucial for distinguishing authentic ancient signals from modern contamination.

The Kraken family of tools [1, 2] generates highly detailed taxonomic classification outputs, which can be overwhelming and difficult to interpret without appropriate filtering. Proper filtering significantly enhances the clarity of the results, yielding a more coherent and biologically meaningful list of identified organisms. In many cases, the filtered output aligns much better with expectations than the raw, unfiltered results. Kraken2 and KrakenUniq offer several key metrics that can be leveraged for filtering. One important metric is the number of reads assigned to each organism, which serves as a proxy for depth of sequencing coverage. Another crucial metric is the number of unique *k*-mers (number of distinct minimizers in Kraken2) associated with an organism, which acts as a proxy for breadth of sequencing coverage. The latter is particularly important, as it helps eliminate false-positive classifications caused by the miss-assignment of reads to highly conserved or convergently evolved genomic regions. This issue has been extensively discussed in previous studies [1, 8], which highlight the risk of erroneous taxonomic identifications arising from sequence similarity in non-specific genomic regions. It was shown in the original KrakenUniq publication [1] that “for the discovery of pathogens in human patients … a unique *k*-mer count threshold of 1000 eliminated many background identifications”. In addition, the thresholds of 1000 unique *k*-mers (breadth of overage) and 200 assigned reads (depth of coverage) were elaborated in [8] and used by default in the ancient metagenomic profiling software aMeta.

As an alternative filtering strategy, E-value, which represents the combination of KrakenUniq filters (number of unique *k*-mers, number of assigned reads and *k*-mer coverage), was previously suggested for metagenomic taxonomic classification by M. Guellil et al. [9] with the optimal threshold varying between 0.001 and 0.1 depending on the dataset. The equation for E-value was later modified by M. Borry [10], by embracing the *k*-mer coverage into the double-exponential function, for a better distinction between true- and false-positive taxonomic assignments by KrakenUniq.

In this study, I perform a comprehensive benchmarking of different filtering approaches using two simulated ancient microbial and one ancient environmental DNA datasets. I compare the suggested in [1, 8] filtering thresholds with the alternative filtering approaches [9, 10] and demonstrate that filtering with respect to the number of unique *k*-mers alone provides the optimal balance between sensitivity and specificity of organism detection in typical metagenomic samples, which is not outperformed by alternative more complex filtering approaches. I also suggest a simple scaling law for extrapolating the suggested filtering threshold (of 1000 for the number of unique *k*-mers) to deeply sequenced metagenomic samples.

## Methods

To benchmark different filtering approaches, I used three ancient metagenomic datasets, simulated using the gargammel tool [11]: (1) a regular microbial dataset, (2) a pathogen-enriched microbial dataset, and (3) an environmental (sedimentary) ancient DNA dataset. The first two datasets were adopted from [8] and each consisted of 10 metagenomic samples (500,000 ancient and 500,000 modern DNA fragments simulated in each sample) with human as the host, mimicking the typical microbial composition observed in ancient Scandinavian human samples [12]. The regular microbial dataset contained 35 microbial species (18 ancient and 17 modern) with microbial DNA percentage varied between 30% and 70%, while the pathogen-enriched dataset included 13 microbial species (9 ancient pathogens and 4 modern contaminants) with microbial reads occupying at most 30% of all reads. Full lists of included microbial species as well ass the details of simulations are available in [8].

Two ancient environmental (sedimentary) DNA (aeDNA) samples were simulated mimicking two opposite situations: (1) comparable DNA contribution from multiple organisms (vertebrates, invertebrates, plants and microbes), and (2) a single organism (presumed mammoth) dominated the DNA composition. In the first case, sixteen organisms across the tree of life were included: human (*Homo sapiens*), African elephant (*Loxodonta africana*), vesper bat (*Pipistrellus kuhlii*), wild boar (*Sus scrofa*), red junglefowl (*Gallus gallus*), saltwater crocodile (*Crocodylus porosus*), brown trout (*Salmo trutta*), thale cress (*Arabidopsis thaliana*), common sunflower (*Helianthus annuus*), black cottonwood (*Populus trichocarpa*), Australian freshwater crayfish (*Cherax quadricarinatus*), East Asian common octopus (*Octopus sinensis*), Old World swallowtail (*Papilio machaon*), and three microbial organisms: *Clostridium botulinum*, *Streptosporangium roseum* and *Yersinia pestis*. In total, 252,498 ancient reads were simulated with 6.25% of reads originating from each of the sixteen species. The second simulated aeDNA sample contained 255,045 ancient reads with 30% being human (*Homo sapiens*), 7% microbial (*Clostridium botulinum*), and 63% of reads coming from African elephant (*Loxodonta africana*). This sample is assumed to mimic the situation of a very contaminated or degraded mammoth sample.

Ancient reads for all samples were simulated with deamination following Briggs parameters [13, 14] in gargammel [11]: *-damage 0.03,0.4,0.01,0.3*. The simulated ancient reads were fragmented and followed a log-normal distribution with the following parameters *--loc 3.7424069808 --scale 0.2795148843*. These parameters were empirically determined from the *Y. pestis* reads in [12]. Illumina sequencing errors were added with the ART module of gargammel [11] to both modern and ancient reads. In addition, Illumina universal sequencing adapters were used, which resulted in 125 bp long paired-end reads.

Simulated ancient metagenomic reads were profiled by KrakenUniq and Kraken2 which employ a Lowest Common Ancestor (LCA) algorithm to handle DNA reads that map equally to multiple organisms. The databases for KrakenUniq and Kraken2 were built using the non-redundant NCBI NT database, which includes reference genomes from microbes, vertebrates, invertebrates, and plants. The database for both tools included identical reference genomes for consistency. The report outputs of KrakenUniq and Kraken2 were filtered using custom scripts, and an extensive benchmarking was conducted across a wide range of filtering thresholds. Figure 1 depicts heatmaps of simulated ground truth across the three datasets as well as the main steps of the benchmarking workflow used in this study.

**Figure 1.**
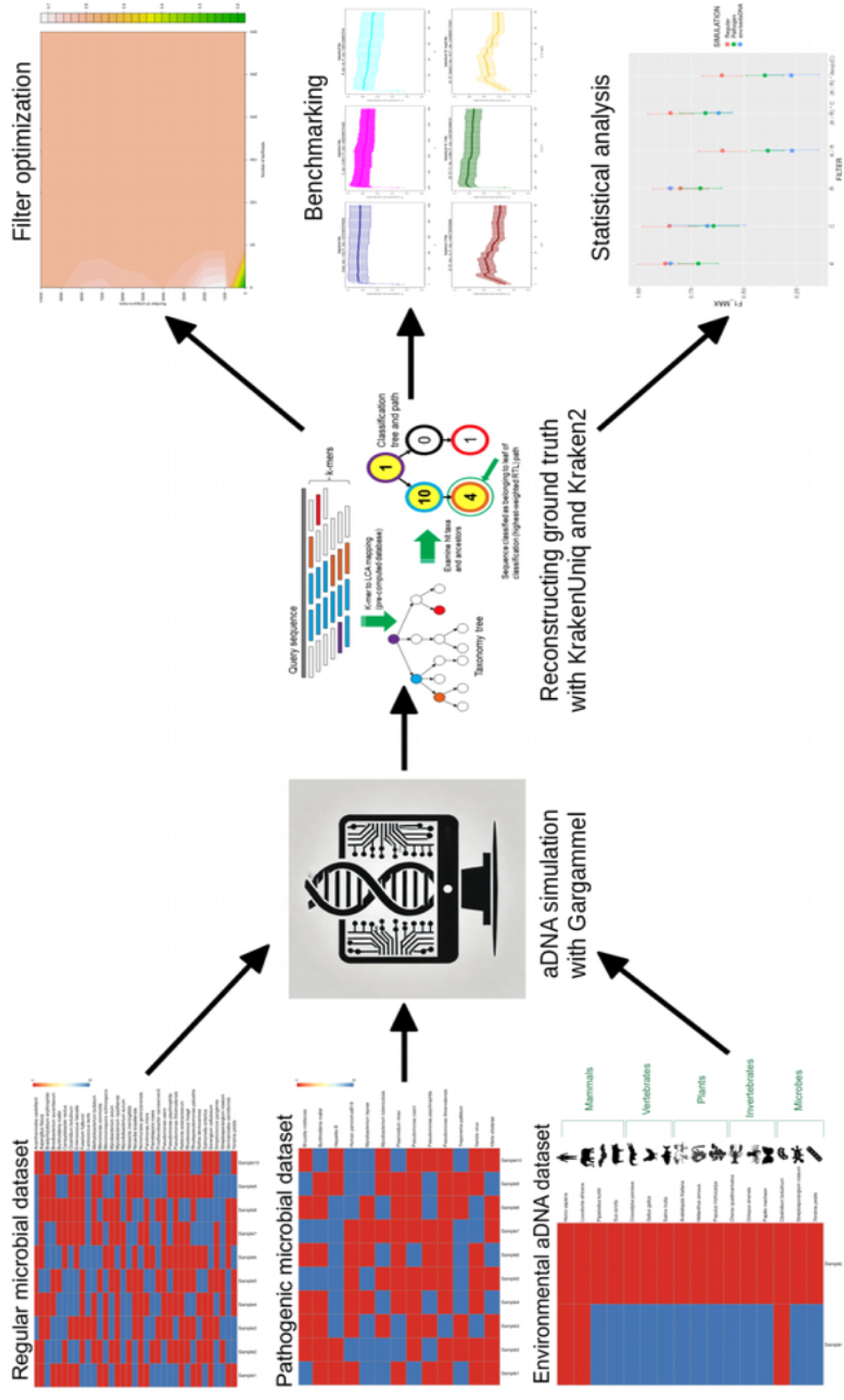
Overview of workflow used for benchmarking different filtering strategies. Heatmaps of the simulated data: red – organism present in the sample, blue – organism absent in the sample.

## Results

To optimize the ground truth reconstruction by KrakenUniq, I explored possible thresholds for the depth (number of assigned reads) and breadth (number of unique *k*-mers) of coverage filters, evaluating their impact on the F1-score—a metric balancing sensitivity and specificity of ground truth reconstruction. The 2D heatmap averaged across 10 simulated regular microbial samples, illustrated in Figure 2, revealed that the number of unique *k*-mers alone was sufficient to achieve the highest F1-score. Specifically, a threshold of at least 1,000 unique k-mers provided optimal filtering, aligning with previously recommended values [1, 8] for conservative microbiome profiling. The corresponding heatmaps computed for the microbial pathogen-enriched dataset and environmental / sedimentary aDNA dataset confirmed the threshold of 1,000 unique *k*-mers as nearly optimal for reconstructing the simulated ground truth, Supplementary Figure 1.

**Figure 2.**
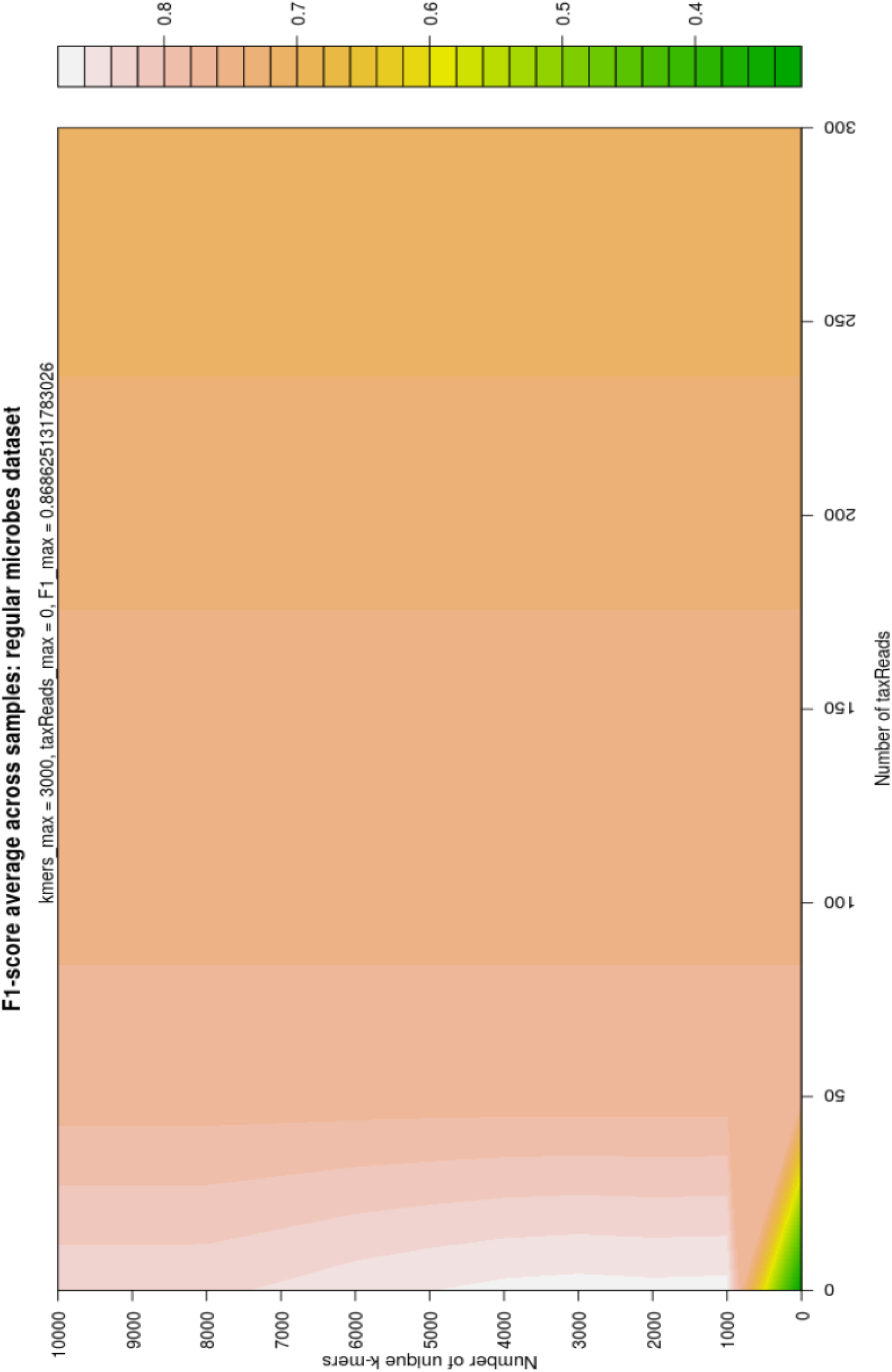
Heatmap F1-score ground truth reconstruction of regular microbial dataset for different values of KrakenUniq depth (number of assigned reads) and breadth of coverage (number of unique *k*-mers) filters.

I further examined how individual filtering metrics affect the accuracy of taxonomic assignment. To achieve this, I explored a wide range of threshold values for six different filters: the number of unique *k*-mers (K filter), the number of assigned reads (R filter), the coverage of unique *k*-mers (C filter), the simple ratio of unique *k*-mers to assigned reads (K/R filter), the E-value-based filter [9] ((K/R) * C filter), and a modified E-value filter according to [10] ((K/R) * dexp(C) filter). Across all three simulated datasets and filtering strategies, the F1-score exhibited a sharp increase, reaching a peak before a gradual decline (Figure 3, Supplementary Figures 2–3). This pattern suggests that a conservative approach with relatively stringent filtering thresholds is consistently more beneficial than a permissive approach. This is explained by the fact that while less stringent filtering may improve sensitivity in detecting organisms within metagenomic samples, it also leads to a high number of false-positive identifications.

**Figure 3.**
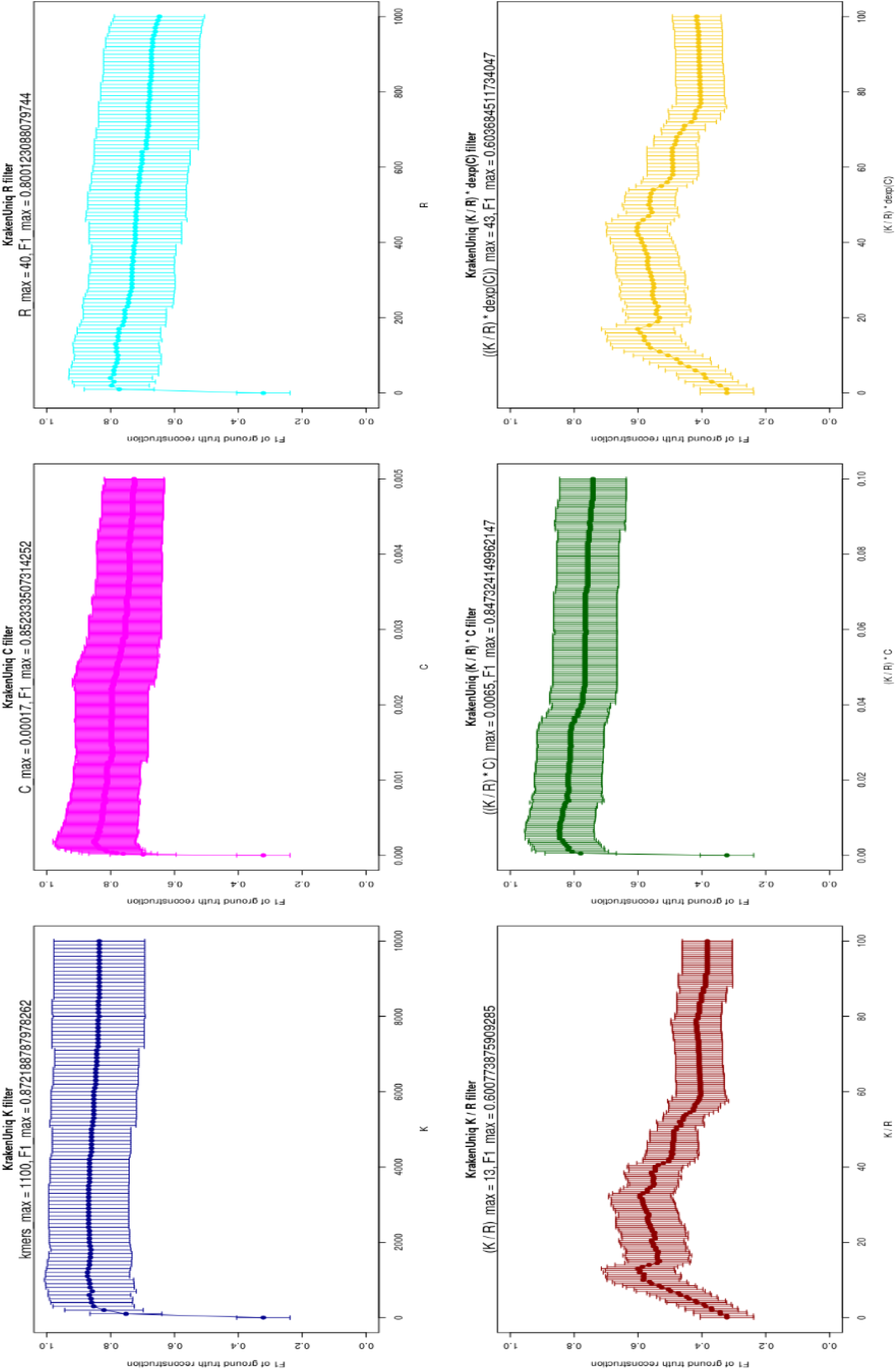
Comparison of different filtering approaches in terms of F1-score for simulated regular microbial dataset. The confidence intervals were computed by averaging across 10 samples.

The optimal number of unique *k*-mers was approximately 1,000, whereas the optimal threshold for assigned reads varied between 40 and 360, depending on the dataset. This aligns with previous findings [8], which indicate that ∼50 assigned reads serve as an absolute minimum for balancing sensitivity and specificity. However, for ancient DNA (aDNA) studies, slightly higher thresholds (∼200 assigned reads) are generally recommended [8]. This is because obtaining a smooth deamination / damage profile with tools like mapDamage [13] becomes challenging when fewer than ∼200 reads are available [8]. Among all the tested filters, the K-filter yielded the highest F1-score across all datasets: 0.87 for the regular microbiome dataset, 0.72 for the pathogen-enriched dataset, and 0.85 for the environmental DNA dataset. In contrast, the K/R filter and the modified E-value filter [10] consistently produced the lowest F1-scores across all three datasets. This trend is clearly illustrated in Figure 4, where I present the maximum F1-scores along with confidence intervals, averaged across all samples in each dataset. While the individual K, C, and R filters performed similarly in terms of F1-score (with the K-filter demonstrating a slight advantage), the combinations of different filters (K/R, (K/R) * C, and (K/R) * dexp(C)) performed significantly worse (Mann-Whittney U test p=0.003), particularly for the environmental DNA dataset. However, the E-value filter [9], defined as (K/R) * C, performed nearly as well as the individual K, R, and C filters for the regular and pathogen-enriched datasets. It is also important to note that although the K, R, and C filters achieve comparable accuracy in reconstructing the ground truth, the E-value filter is significantly less interpretable and more challenging to understand intuitively. Based on this comparison, I conclude that simple filtering based on unique *k*-mers (K-filter) alone is sufficient for accurate reconstruction of the simulated ground truth, and combining multiple filters does not lead to a significant improvement in performance.

**Figure 4.**
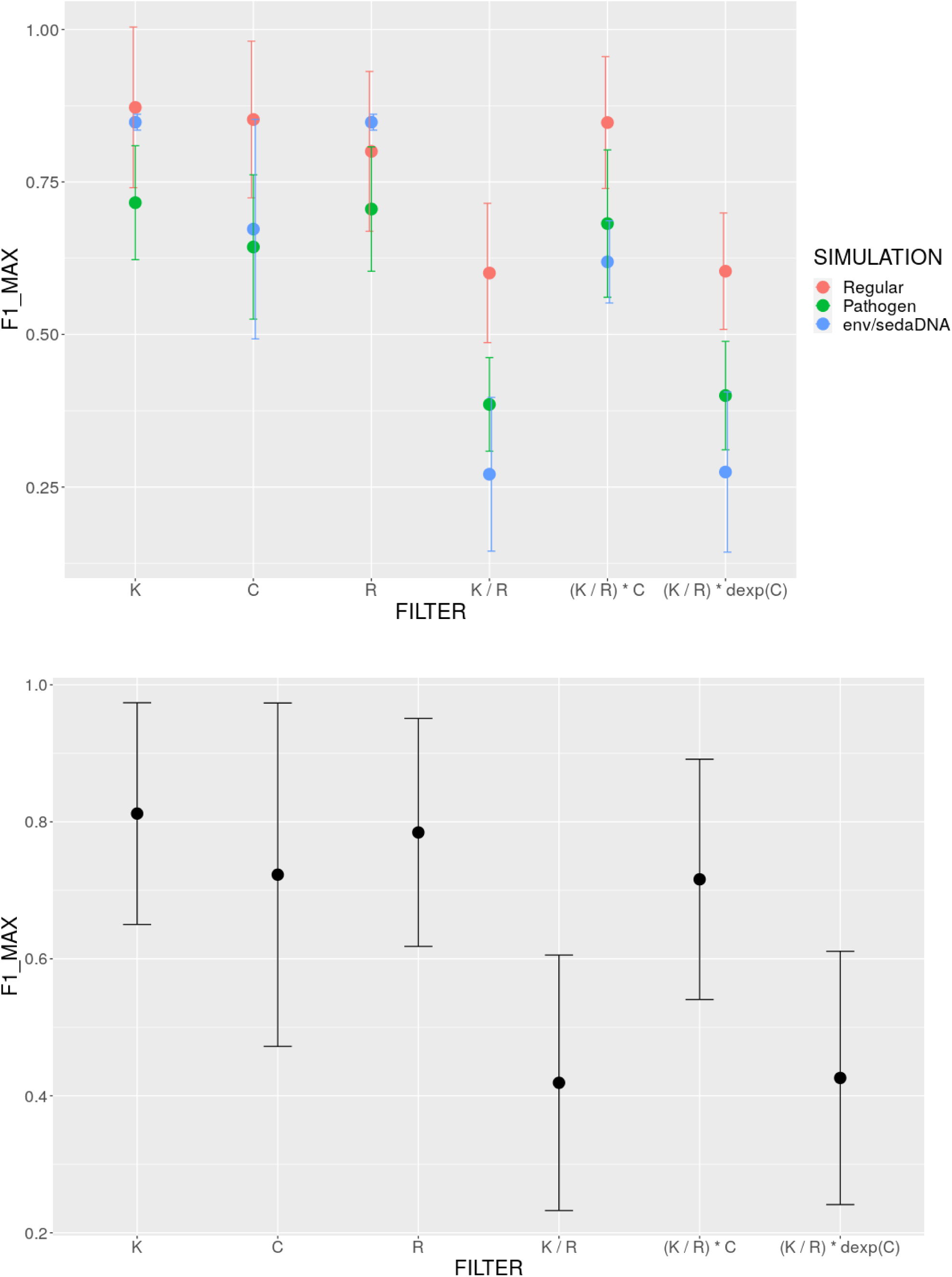
F1-score comparison of different filtering approaches: a) three simulated datasets (regular microbial, pathogen-enriched, environmental ancient DNA) separately, b) average across datasets.

To examine the relationships between different KrakenUniq filtering metrics, I generated a Spearman correlation heatmap for each dataset (Figure 5 and Supplementary Figures 4–5). Overall, the heatmaps reveal a high degree of correlation among the metrics provided by KrakenUniq. In particular, the percentage of reads, total reads, taxReads, and the number of unique k-mers exhibit strong correlations, with Spearman’s rho values ranging from approximately 0.7 to 0.9. This finding helps explain why filtering based on either the number of unique *k*-mers or the number of assigned reads resulted in comparable F1-scores for reconstructing the ground truth (Figures 3 and 4). In contrast, coverage and, more notably, the number of duplicates show weaker correlations with the other metrics, with Spearman’s rho values ranging from approximately 0.3 to 0.7.

**Figure 5.**
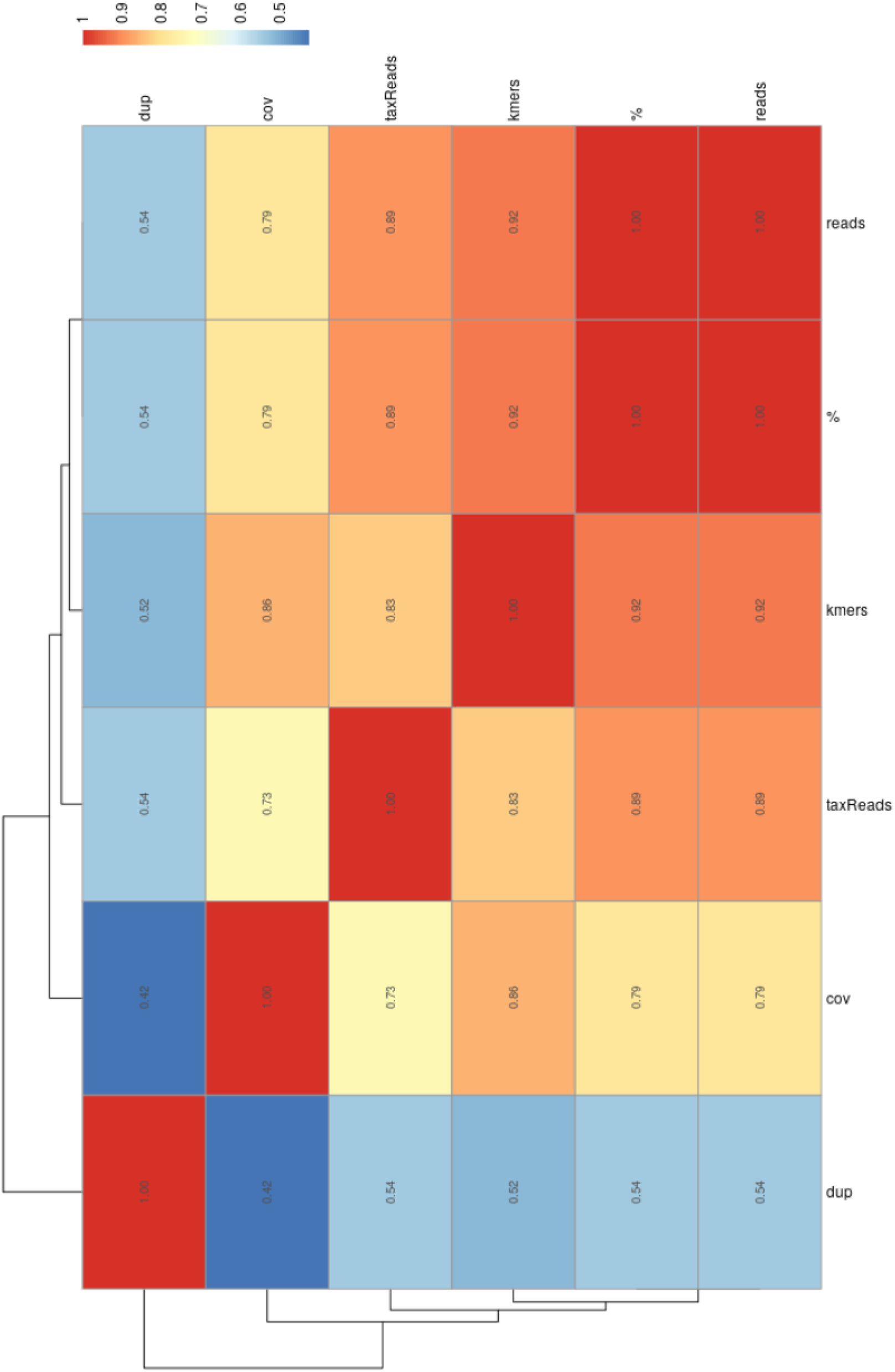
Pairwise Spearman correlation heatmap across KrakenUniq filters for regular microbial dataset averaged across all samples.

Since Kraken2 and KrakenUniq employ different strategies for database construction and use slightly different breadth of coverage filters—KrakenUniq relies on the number of unique *k*-mers, while Kraken2 uses the number of distinct minimizers—I aimed to compare their performances on regular and pathogen-enriched microbial datasets. To ensure a fair comparison, both tools were tested using databases containing identical reference genomes, and exhaustive filtering optimization was applied to their results. The key question was whether, given the most optimal filtering conditions for both KrakenUniq and Kraken2, their accuracy in reconstructing the ground truth would be comparable. Supplementary Figures 6 and 7 show that, although the optimal thresholds for unique *k*-mers and distinct minimizers differ significantly, the resulting F1-scores are largely comparable, with KrakenUniq exhibiting slightly higher accuracy. Interestingly, Kraken2 appears to require an optimal number of distinct minimizers that is at least 2–3 times higher than the optimal number of unique k-mers used by KrakenUniq.

Finally, I aimed at exploring whether deep sequencing of metagenomic samples can affect the number of unique *k*-mers threshold of 1,000 for KrakenUniq. I found that the optimal value of this threshold is significantly correlated (p=7*10^-7^) with sequencing depth and should be adjusted for deeply sequenced metagenomic samples. By determining the optimal number of unique *k*-mers that maximize the F1-score for ground truth reconstruction across 22 simulated samples, I observed that this optimal value increases approximately linearly with sequencing depth. Specifically, the relationship follows the approximate formula: *optimal_n_unique_kmers ∼ 0.002 * seq_depth*. This translates to roughly 200 unique *k*-mers per 100,000 reads, as illustrated in Supplementary Figure 8. This simple scaling rule can be important to keep in mind when working with deeply sequenced metagenomic samples.

## Discussion

Taxonomic classification of metagenomic data remains a cornerstone in profiling both past and modern prokaryotic and eukaryotic environments. The choice of an appropriate taxonomic classification tool is a topic of ongoing debate in the field of ancient metagenomics, particularly as different methods employ distinct algorithms and heuristics. Several benchmarking studies have attempted to compare the performance of various taxonomic profilers, including Kraken2 and KrakenUniq, yet a consistent challenge lies in the fair assessment of these tools due to differences in how they filter their classification outputs [5–7].

In this study, I systematically evaluated different filtering strategies for KrakenUniq and Kraken2 using three simulated datasets: a regular microbial dataset, a pathogen-enriched microbial dataset, and an environmental ancient DNA (aDNA) dataset. My results suggest that filtering based on the number of unique *k*-mers is the most effective strategy for optimizing the accuracy of taxonomic assignment. Unlike other filtering approaches, such as those based on the number of assigned reads or combined filtering metrics, I observed that the unique k-mer filter alone provided the highest F1-scores for ground truth reconstruction. This finding simplifies the filtering process and supports the use of unique *k*-mers as a reliable metric for reducing false-positive assignments in metagenomic studies.

Furthermore, my comparison of Kraken2 and KrakenUniq demonstrates that, although both tools apply different database construction strategy, they yield comparable accuracy when optimized filtering parameters are applied. Although KrakenUniq relies on the number of unique *k*-mers, while Kraken2 utilizes the number of distinct minimizers, my results indicate that, after exhaustive optimization, their performance differences are minimal. This highlights the importance of proper filtering rather than the choice of the classifier itself. Interestingly, Kraken2 required 2-3 times more distinct minimizers to achieve a comparable level of accuracy to KrakenUniq, suggesting that the two classifiers differ in their sensitivity to sequence complexity and taxonomic resolution.

Another key finding of this study is that the optimal number of unique k-mers increases approximately linearly with sequencing depth. Specifically, I derived a simple scaling law indicating that the optimal threshold for unique k-mers is approximately 200 per 100,000 reads. This relationship provides a practical guideline for researchers working with metagenomic datasets of varying sequencing depths, ensuring that filtering thresholds are appropriately adjusted to maintain accuracy without overly conservative exclusion of true positives.

Despite these insights, several limitations should be considered. First, my benchmarking was conducted on simulated datasets, which, while valuable for controlled comparisons, may not capture the full complexity of real-world metagenomic samples, particularly in the presence of sequencing artifacts or uncharacterized microbial diversity. Second, while I identified unique *k*-mers as the most effective filter in my study, future research should explore whether this conclusion holds across different reference databases and taxonomic groups, including non-microbial components of metagenomes.

In conclusion, this study provides strong evidence that filtering by the number of unique *k*-mers is an optimal strategy for taxonomic classification in ancient metagenomics. By applying this approach, researchers can enhance the accuracy of microbial and environmental profiling while minimizing false-positive taxonomic assignments. Additionally, the comparable performance of KrakenUniq and Kraken2 underscores the importance of filtering optimization over classifier selection. These findings contribute to the ongoing refinement of best practices in metagenomic data analysis and offer practical guidelines for filtering strategies that can be adapted to diverse sequencing depths and study designs.

## Acknowledgments

I am financially supported by Knut and Alice Wallenberg Foundation as part of the National Bioinformatics Infrastructure Sweden at SciLifeLab. I am particularly grateful to the ancient metagenomics SPAAM community, and especially Maxim Borry, Zoe Pochon and Meriam Guellil for fruitful discussions and inspiration for this article.

## Data and Code Availability

The simulated microbial datasets, can be accessed via the SciLifeLab Figshare at https://doi.org/10.17044/scilifelab.21261405 and https://doi.org/10.17044/scilifelab.24211584. The codes used for simulating the ancient metagenomic samples are available in the GitHub repository: https://github.com/NikolayOskolkov/aMeta, and Zenodo https://doi.org/10.5281/zenodo.8130819. The pre-built KrakenUniq database based on reference sequences from the non-redundant NCBI NT is available at https://doi.org/10.17044/scilifelab.20205504. The simulated ancient environmental DNA samples, Kraken output files for all three datasets as well as scripts used for the simulation and analysis are available at https://github.com/NikolayOskolkov/KrakenFilterManuscript.

## Supplementary Figures

**Supplementary Figure 1.**
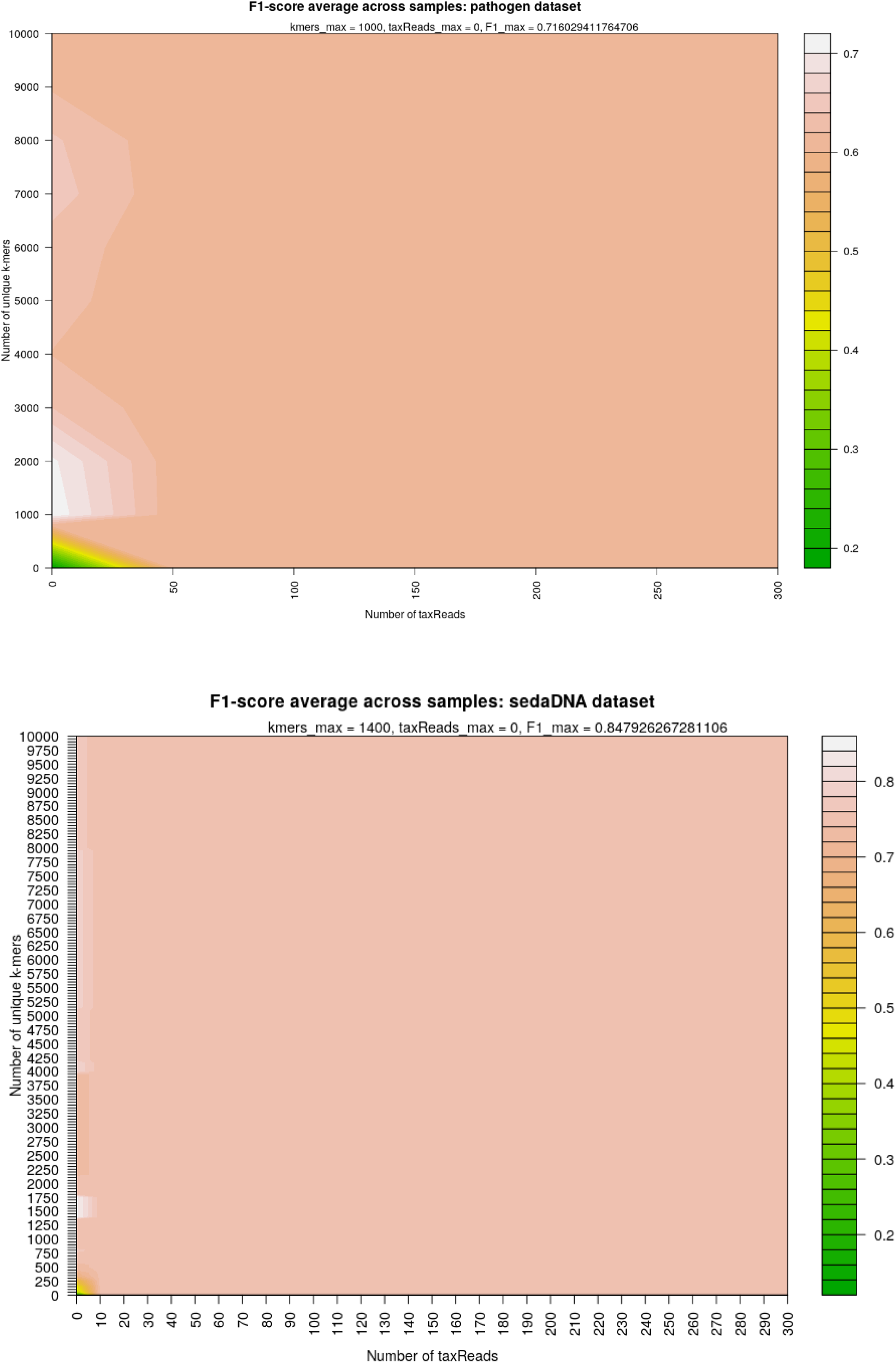
Heatmap of optimal ground truth reconstruction F1-score of microbial pathogen-enriched and environmental / sedimentary aDNA datasets.

**Supplementary Figure 2.**
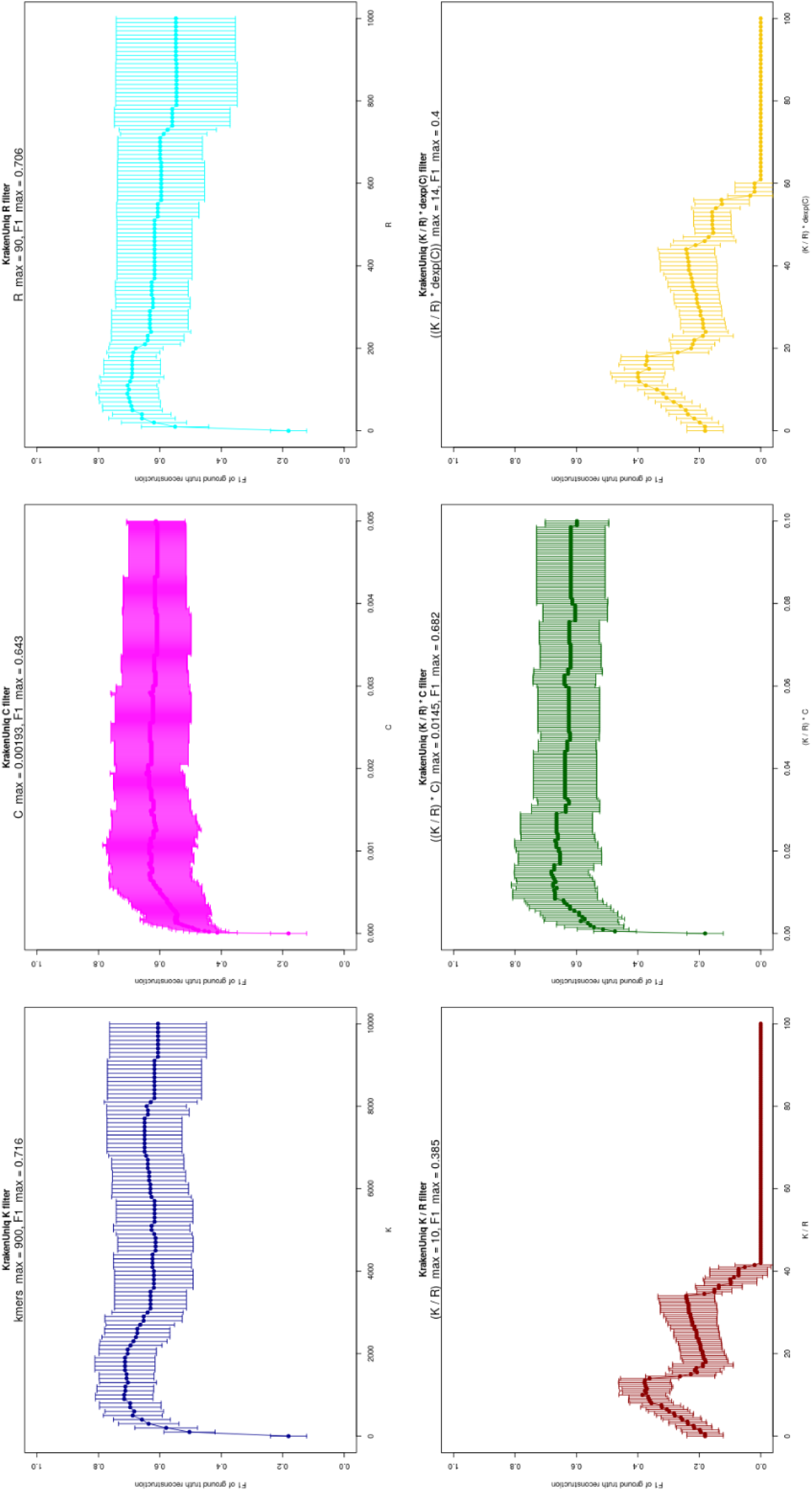
Comparison of different filtering approaches in terms of F1 score for simulated microbial pathogen-enriched dataset.

**Supplementary Figure 3.**
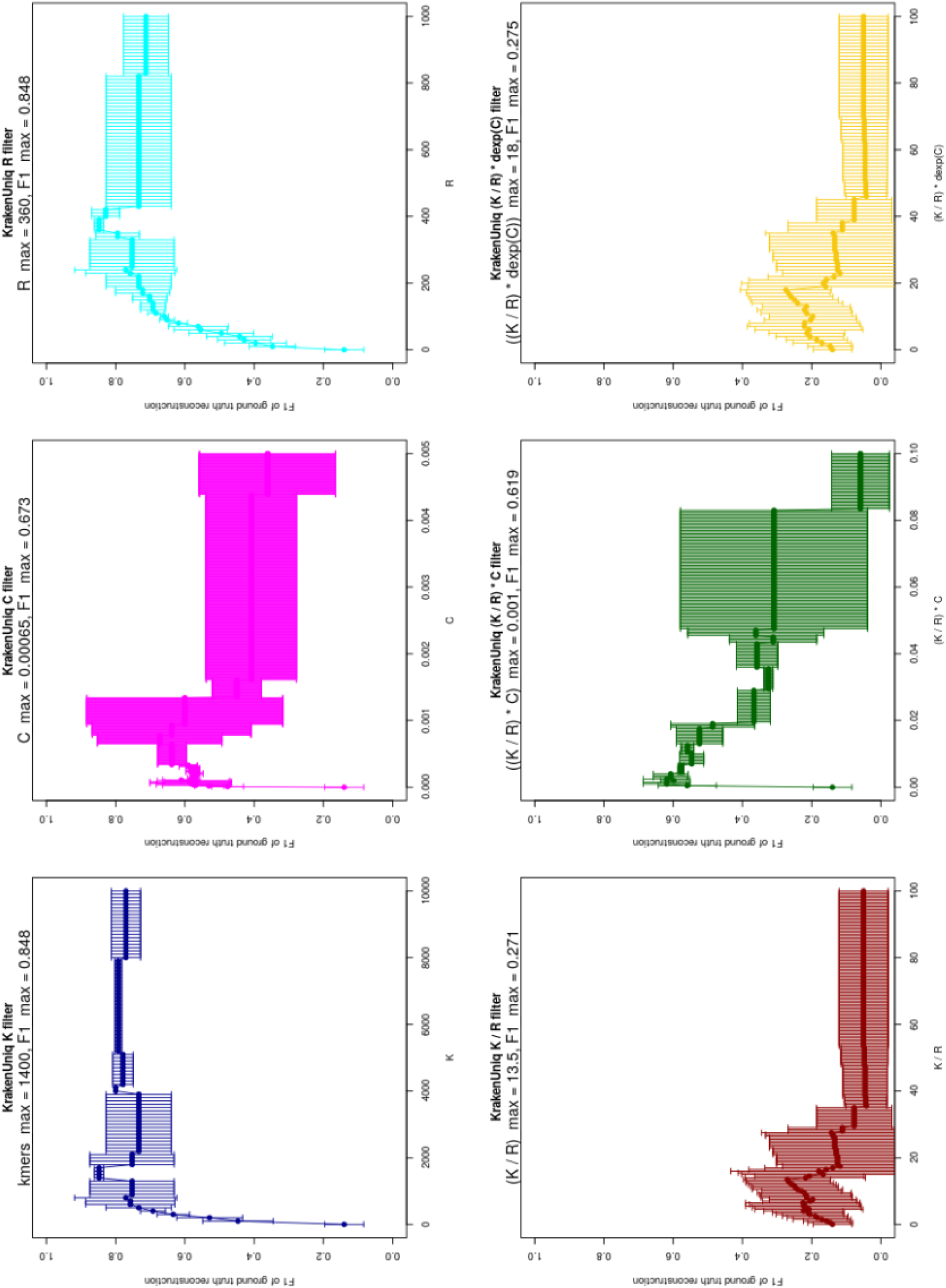
Comparison of different filtering approaches in terms of F1 score for simulated environmental / sedimentary DNA dataset.

**Supplementary Figure 4.**
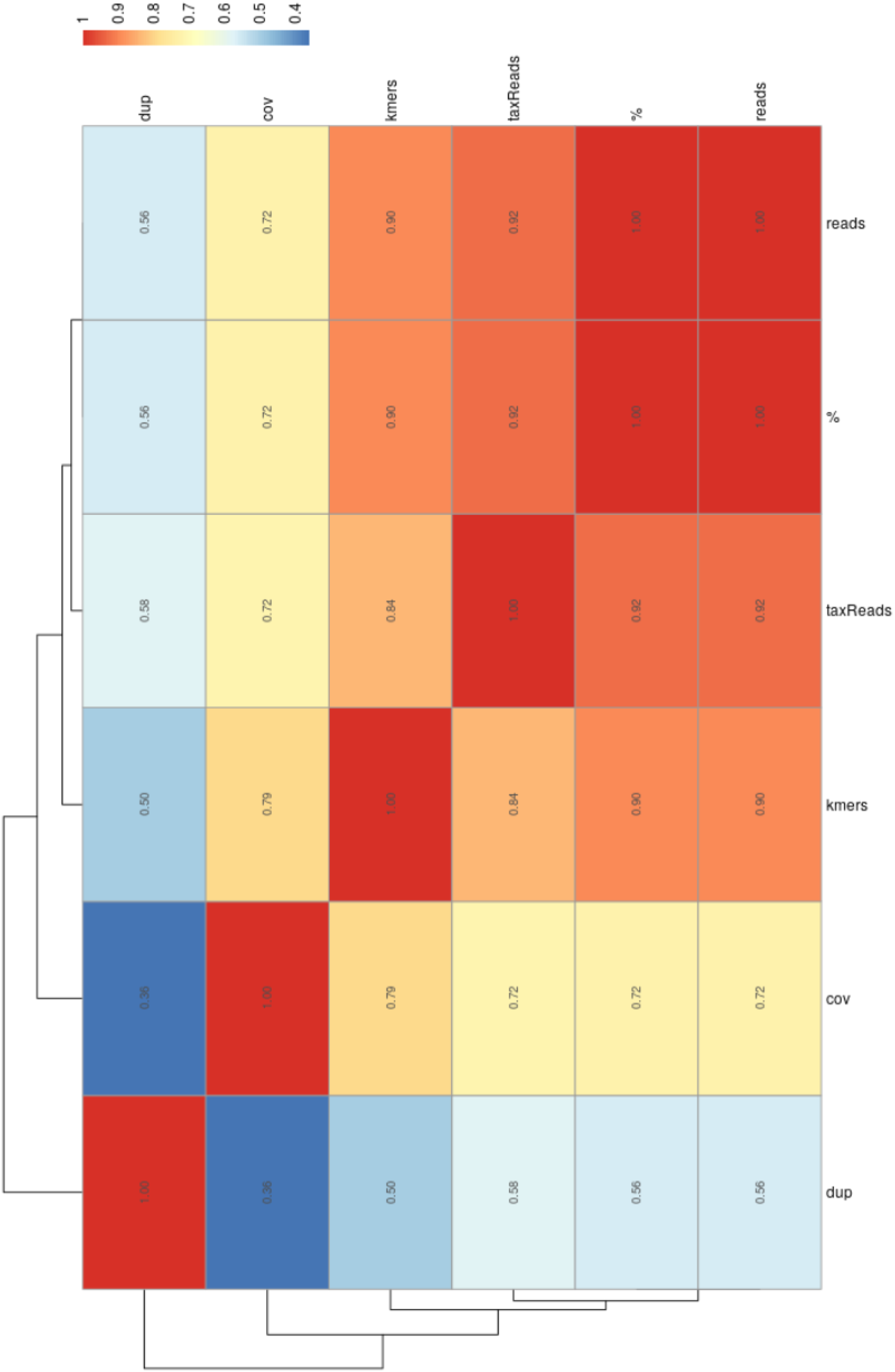
Pairwise Spearman correlation heatmap across KrakenUniq filters for pathogen-enriched microbial dataset averaged across all samples.

**Supplementary Figure 5.**
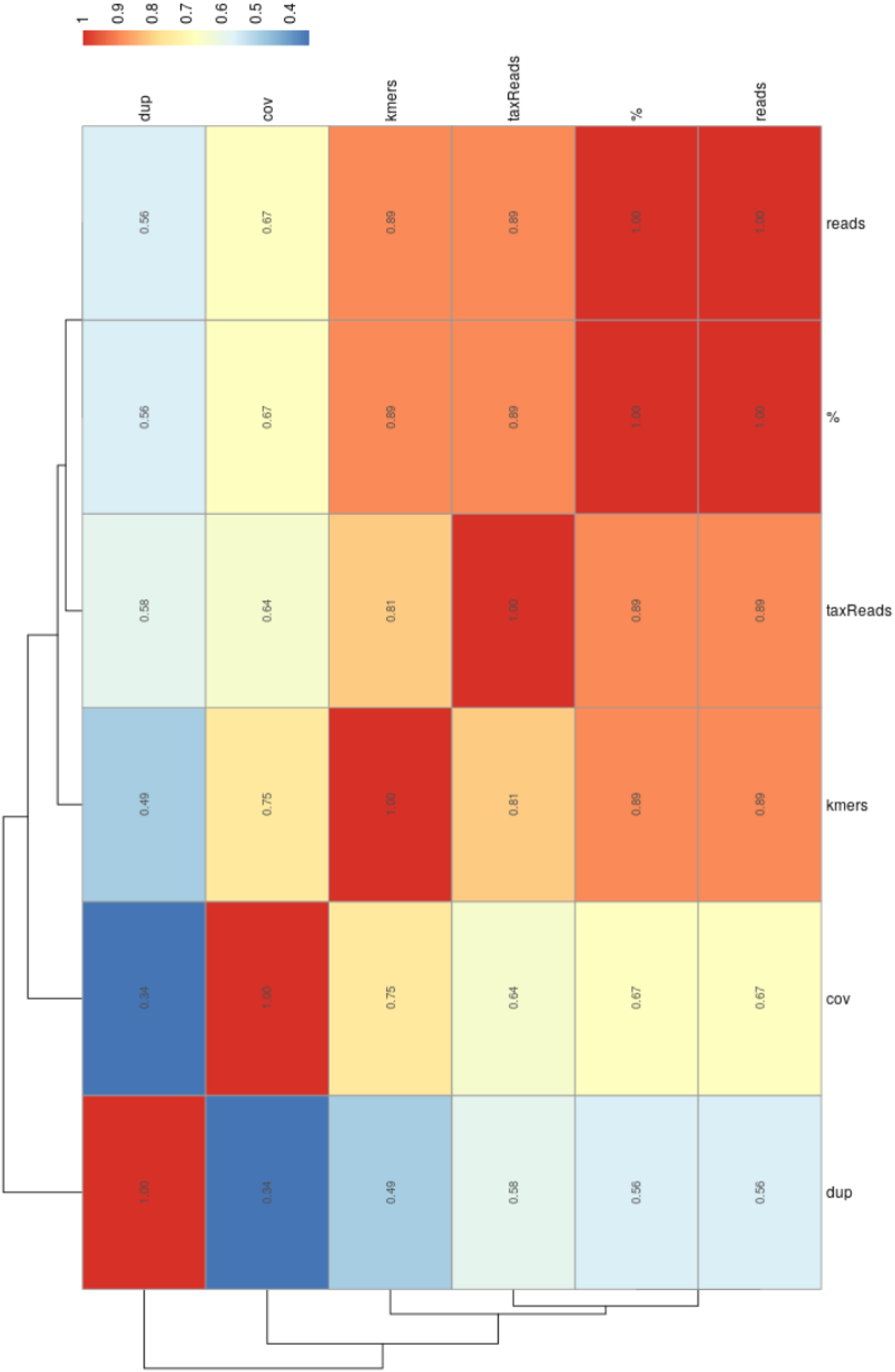
Pairwise Spearman correlation heatmap across KrakenUniq filters for environmental DNA dataset averaged across all samples.

**Supplementary Figure 6.**
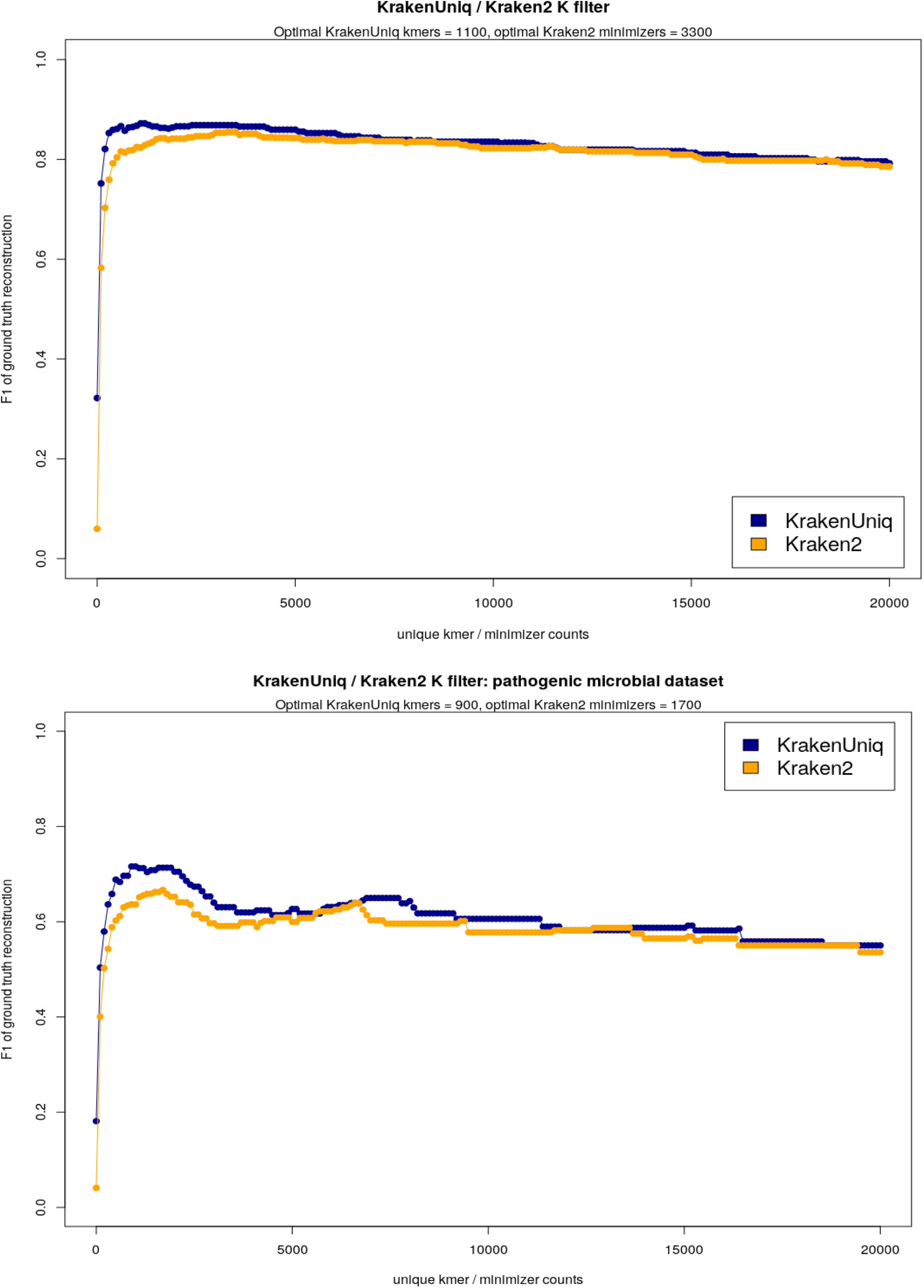
Comparison of KrakenUniq vs. Kraken2 performance on a) regular microbial, and b) pathogen-enriched microbial datasets. In both case, the plots correspond to optimal filtering strategies applied to KrakenUniq and Kraken2 outputs.

**Supplementary Figure 7.**
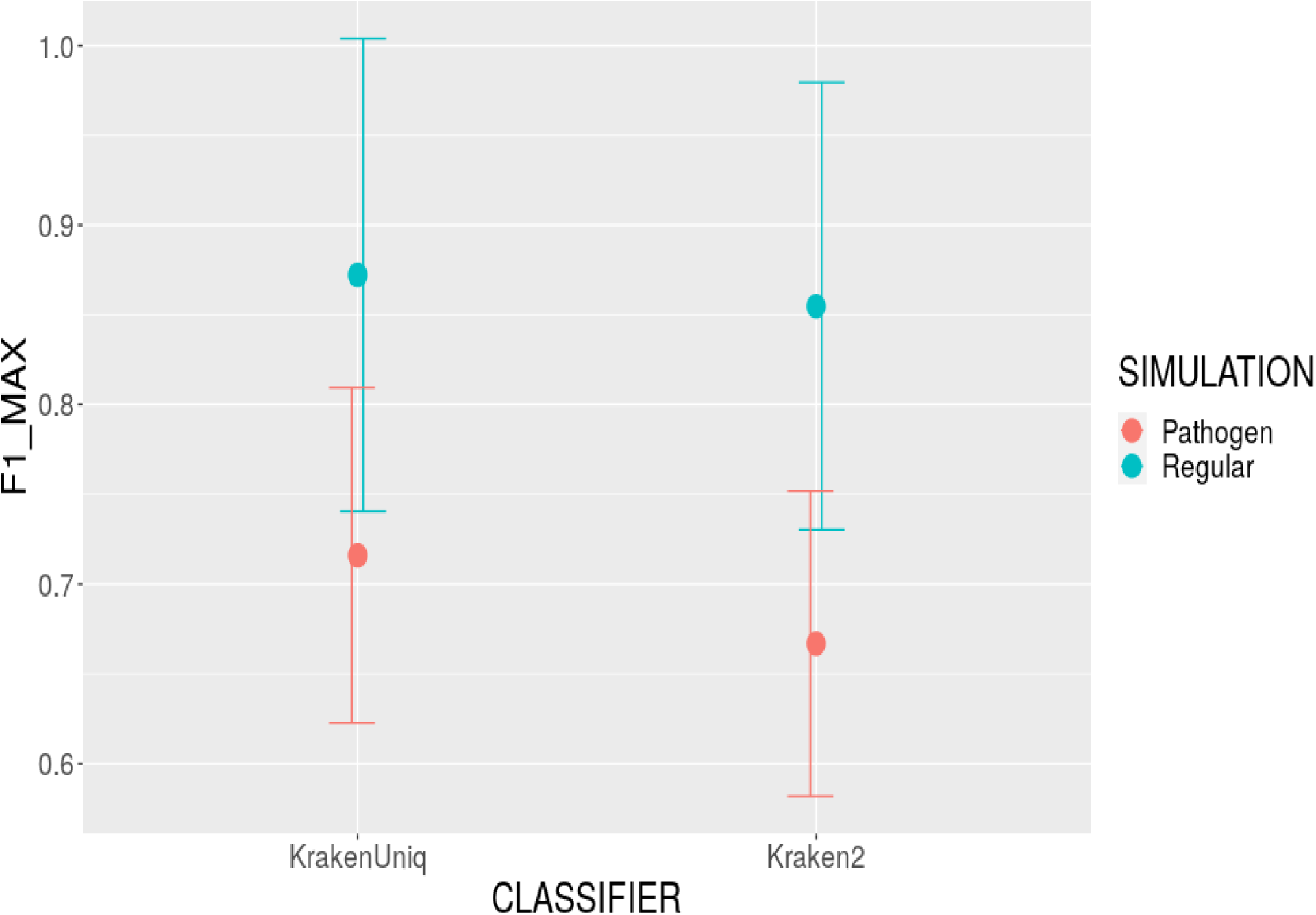
Comparison of optimal F1-scores for KrakenUniq vs. Kraken2 performance on a) regular microbial, and b) pathogen-enriched microbial datasets. The difference in F1-score is not statistically significant.

**Supplementary Figure 8.**
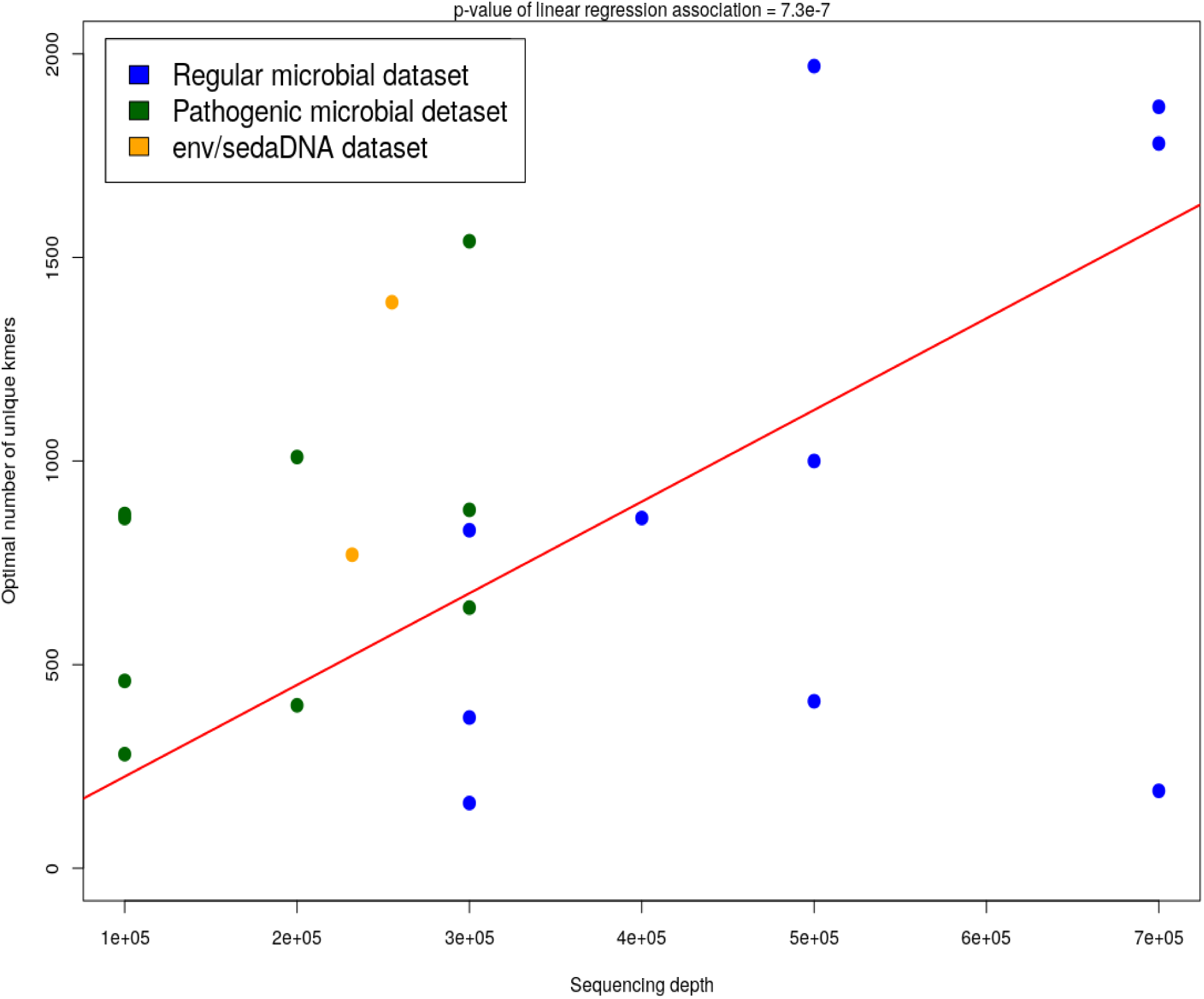
Optimal number of unique *k*-mers grows approximately linearly with the sequencing depth following approximate law optimal_n_unique_kmers ∼ 0.002 * seq_depth.

## Notes

### Competing Interest Statement

The authors have declared no competing interest.

